# Clonal Hematopoiesis Mutations Increase Risk of Alzheimer’s Disease with *APOE* ε3/ε3 Genotype

**DOI:** 10.1101/2025.05.19.654981

**Authors:** Jaejoon Choi, Kyung Sun Park, Yann Le Guen, Jong-Ho Park, Zinan Zhou, Liz Enyenihi, Ila Rosen, Yoon-Ho Choi, Christopher A. Walsh, Jong-Won Kim, August Yue Huang, Eunjung Alice Lee

## Abstract

Clonal hematopoiesis of indeterminate potential (CHIP) represents clonal expansion of blood cells, and increases the risk of hematological malignancies and cardiovascular disorders. Recent studies have studied CHIP mutations in individuals with Alzheimer’s disease (AD), but it is unclear whether their role in AD pathogenesis is protective, detrimental, or neutral. In this study, we used molecular-barcoded deep gene panel sequencing (∼400X) to examine CHIP mutations in 298 blood samples from AD and neurotypical individuals 60 years and older. The AD patients exhibited a significantly higher burden of CHIP mutations compared to the age-matched controls (p < 2e-7, odds ratio (OR) = 2.89), particularly in low-frequency variants often not captured by standard whole exome or whole genome sequencing (WGS). This increase was driven by individuals with the *APOE* ε3/ε3 genotype and absent in ε4 carriers. Analysis of an independent dataset from the Alzheimer’s Disease Sequencing Project (ADSP), comprised of WGS data from ∼30,000 individuals, confirmed increased CHIP mutations in AD versus control (p < 0.02, OR = 1.32), again driven by individuals with *APOE* ε3/ε3 genotype. CHIP mutations in AD patients also showed stronger positive selection than in controls. Our results indicate that AD patients show significantly more CHIP mutations in their blood than controls, involving more than one third of AD patients, and contributing to AD risk through a mechanism independent of *APOE* ε4.

## Main Text

Alzheimer’s Disease (AD) is a progressive neurodegenerative disorder characterized by cognitive decline and memory loss, with advanced age and the *APOE* ε4 allele as established risk factors^1,2^. Somatic mutations are DNA changes acquired postzygotically that accumulate with age in various cell types and have been reported to contribute to a variety of age-associated diseases, including AD^3–5^. Clonal hematopoiesis of indeterminate potential (CHIP) is the clonal expansion of hematopoietic stem cells characterized by the acquisition of somatic mutations in cancer driver genes^6–8^. Individuals with CHIP mutations are commonly asymptomatic; however, they have an elevated risk of developing leukemia due to the increased proliferative potential of mutant blood cells^7^. Additionally, CHIP has been found to promote cardiovascular disease^9^, likely by promoting a pro-inflammatory environment that affects surrounding non-mutant cells.

Recent studies have reported an increased burden of somatic cancer driver mutations, notably in driver genes for CHIP, in the brains of AD patients, particularly in microglia, the primary resident immune cells of the brain, and identical mutations are frequently found in the peripheral blood of the same donors^10,11^. However, the role of CHIP mutations in the blood in AD pathogenesis remains controversial. While some studies suggest a potential positive correlation between CHIP mutations and AD risk^12^, others have reported conflicting results, such as no correlation^13,14^ or even a protective effect^15^. These discrepancies may reflect factors such as limited sensitivity in detecting somatic mutations due to low sequencing depth (usually < 30X for WGS and 50–70X for exome-wide sequencing) and/or inadequate control of confounding variables.

To address these limitations, we first conducted a comprehensive profiling of somatic mutations in 149 cancer driver genes in the peripheral blood of 298 individuals: 108 AD patients, 154 age-matched controls and 36 centenarians without AD, utilizing molecule-barcoded targeted panel sequencing^10^ (Fig. 1A). To minimize confounding effects, we included only individuals with *APOE* genotypes ε3/ε3 or ε3/ε4. As expected, the AD group had a slightly higher proportion of *APOE* ε3/ε4 carriers than the controls, consistent with the well-established role of the *APOE* ε4 allele as a risk factor for AD. Our approach achieved a high and comparable sequencing depth of ∼400X in both the AD and control groups after collapsing reads using unique molecular identifiers (UMIs) (Extended Data Fig. 1). This UMI-based consensus calling eliminated sequencing errors, enabling the accurate identification of somatic mutations even at variant allele fractions (VAFs) below 0.5%, with a validation rate of 91.5% confirmed by amplicon sequencing (Extended Data Fig. 2).

**Figure 1.**
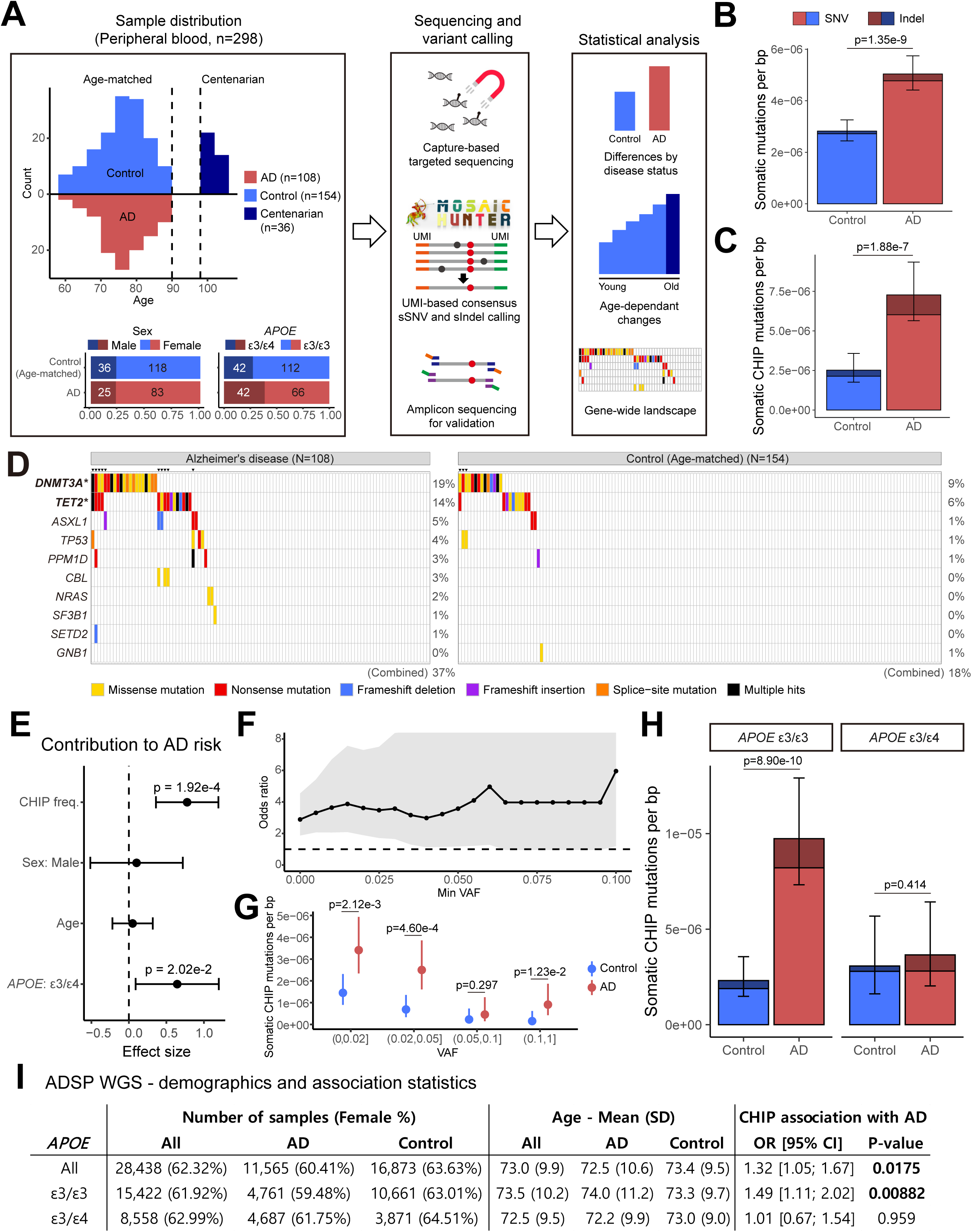
Elevated burden of somatic cancer driver mutations in AD blood samples compared to controls. **A.** Profiling of somatic mutations in the blood samples of 108 AD patients and 190 ndividuals (including 154 age-matched controls). AD and age-matched controls had similar sex hile AD individuals had a slightly higher proportion of *APOE* ε3/ε4 genotypes. **B-C.** AD patients significantly more somatic mutations than age-matched controls, whether considering **(B)** all or **(C)** only mutations previously reported to drive CHIP. Error bars represent 95% confidence (CIs). **D.** Top 10 recurrently mutated genes in AD and age-matched control blood samples hit by utations. Asterisks denote genes with significantly more CHIP mutations in AD patients than in ched controls (p < 0.05, one-sided proportion test). Triangles highlight individuals who carry CHIP in multiple genes. **E.** The number of CHIP mutations was significantly associated with AD risk, ntrolling for potential confounding factors including *APOE* ε3/ε4 genotypes, which contribute to AD pendently. Error bars represent 95% CIs. **F-G.** Consistently higher burden of CHIP mutations was in AD patients compared to age-matched controls, with stronger signals driven by mutations with AF. Error bars represent 95% CIs. **H.** The increase of CHIP mutations in AD patients was nantly observed in *APOE* ε3/ε3 individuals, but not in ε3/ε4 individuals. Error bars represent 95% n independent dataset with a larger cohort, ADSP, also showed an increase of CHIP mutations in nts with *APOE* ε3/ε3. Significant p-values are highlighted in bold.

Firstly, we compared the somatic mutation burden between 108 AD patients and 154 age-matched controls (For the remainder of this study, the term ‘control(s)’ refers specifically to these age-matched controls) with a comparable sex distribution (Fig. 1A). We found that AD blood samples harbored a significantly higher burden of somatic mutations across all targeted cancer driver genes (p = 1.35×10^-9^, odds ratio (OR) = 1.79, one-sided proportion test; Fig. 1B). Since not all somatic mutations in cancer driver genes confer a proliferative advantage, we then restricted our analysis to somatic mutations previously reported in CHIP^15^ and observed an even stronger increase in mutation burden in AD patients (p = 1.88×10^-7^, OR = 2.89; Fig. 1C). Our gene-wide landscape analysis further confirmed this AD enrichment, revealing that 37% of AD patients harbored CHIP mutations in the top ten most frequently mutated genes, compared to only 18% of age-matched controls (Fig. 1D and Extended Data Fig. 3). Notably, two genes (*DNMT3A* and *TET2*) showed significant AD enrichment at the individual gene level, both of which are well-established CHIP-associated genes. Furthermore, we observed a significantly higher frequency of individuals carrying mutations in multiple CHIP genes among AD patients compared to controls (10 of 108 in AD vs 3 of 154 in control, p = 9.30×10^-3^, two-sided Fisher’s exact test), suggesting that these mutations may synergistically contribute to AD risk.

This increased CHIP mutation burden was shown as a significant contributor to AD risk (p = 1.92×10^-4^, OR = 2.18, logistic regression model), independent of potential confounding factors, including sex, age, and *APOE* genotype (Fig. 1E). As our dataset was age- and sex-matched between AD and control groups, these factors did not significantly explain AD status. However, as expected, the *APOE* ε4 genotype was a significant risk factor for AD in our regression model (p = 2.02×10^-2^, OR = 1.90). The observed increase in somatic CHIP mutations in AD patients remained robust across various VAF cutoffs (Fig. 1F). Notably, the differences between AD and controls were more pronounced and statistically significant for variants with lower VAFs (VAF ≤ 0.02: p = 2.12×10^-3^, OR = 2.35, one-sided proportion test; 0.02 < VAF ≤ 0.05: p = 4.60×10^-4^, OR = 3.64; Fig. 1G), underscoring the contribution of these low-VAF mutations to AD risk and highlighting the importance of deep sequencing for their detection. Interestingly, our analysis revealed that the increase in CHIP mutations was primarily driven by individuals with the *APOE* ε3/ε3 genotype (p = 8.90×10^-10^, OR = 4.21, one-sided proportion test; Fig. 1H), whereas no significant association was observed in *APOE* ε3/ε4 carriers (p = 0.414, OR = 1.19).

To further validate our findings with a larger sample size, we performed a similar analysis on an independent dataset from the Alzheimer’s Disease Sequencing Project (ADSP)^16^ (Fig. 1I). This analysis included a large cohort of 11,565 AD patients and 16,873 age-matched healthy controls from the ADSP WGS data. We identified individuals carrying CHIP mutations by extracting and annotating variants in known CHIP-related genes^15^, applying stringent filtering criteria to minimize false positives (see Methods). Using a generalized linear model while controlling for sex, age-at-visit, *APOE* ε2 and ε4 dosages, and genetic ancestry, we again observed a significantly higher burden of CHIP mutations in AD patients compared to controls (p = 0.0175, OR = 1.32, logistic regression model). Consistent with our panel sequencing results, this association was specifically observed in AD patients with *APOE* ε3/ε3 genotypes (p = 8.82×10^-3^, OR = 1.49), but not in ε3/ε4 carriers (p = 0.959, OR = 1.01). These findings suggest the possibility of distinct AD pathogenic mechanisms associated with different *APOE* genotypes, with CHIP mutations potentially playing a more prominent role in individuals with the *APOE* ε3/ε3 genotypes.

By analyzing a cohort of 190 individuals without diagnosed neurological diseases, including 36 centenarians, our panel sequencing revealed an increase in CHIP mutations with age in neurotypical controls, both in the proportion of mutation carriers (Fig. 2A) and overall mutation burden (Fig. 2B), aligning with previous reports^7,8^. However, CHIP accumulation occurred at a significantly faster rate in AD patients than in controls (p = 1.07×10^-4^, interaction term in linear regression model), leading to more pronounced burden differences in older age groups. The mutation burden observed in centenarians followed the age-related trend established by the control group. This contrasts with the accelerated accumulation observed in AD patients.

**Figure 2.**
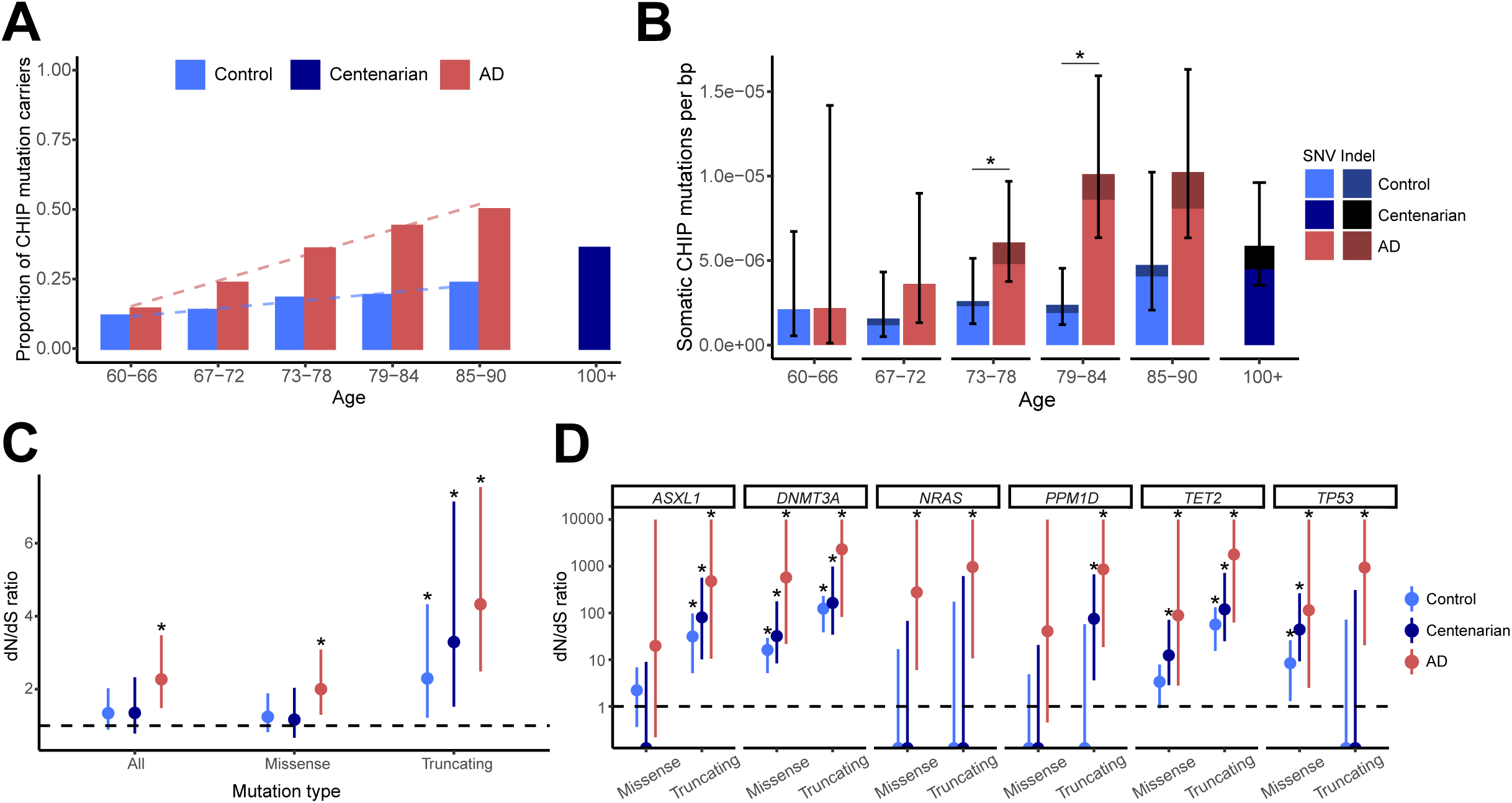
Accelerated somatic CHIP mutation accumulation and stronger positive selection in AD mples. A-B. Somatic CHIP mutations accumulate much faster in AD blood samples than in both in the proportion of CHIP mutation carriers **(A)** and somatic CHIP mutation burdens **(B)**. ans (age of 100 or higher) showed expected mutation frequencies consistent with an extension of group. Asterisks denote age groups that have significantly higher mutation burden in AD to controls (p < 0.05, one-sided proportion test). Error bars represent 95% CIs. **C-D**. Somatic in AD samples showed higher dN/dS ratios across all panel genes **(C)** and in the top six mutated genes **(D)**, which indicate stronger positive selection in AD blood. Asterisks denote io significantly higher (p < 0.05, hypothesis test using confidence interval) than dN/dS=1 (black e). Error bars represent 95% CIs.

To investigate the driving force behind the accumulation of somatic mutations in AD patients, we assessed the selective pressures acting on these mutations by analyzing the ratio of non-synonymous to synonymous substitutions (dN/dS) (Extended Data Fig. 4). We found that somatic mutations in AD samples exhibited a higher dN/dS ratio compared to age-matched controls and centenarians (Fig. 2C), indicating stronger positive selection acting on somatic mutations in AD patients than in controls. Notably, the top six genes showing the strongest positive selection signals in AD (*ASXL1*, *DNMT3A*, *NRAS*, *PPM1D*, *TET2*, and *TP53*; Fig. 2D) are all well-established CHIP-associated genes^6,8,13,15,17,18^, suggesting that these somatic mutations may confer a proliferative advantage to the cells carrying them, and that AD may create an environment that selects for mutant clones. Centenarians displayed an intermediate dN/dS ratio, positioned between the control and AD groups.

In this study, we revealed a contribution of somatic CHIP mutations to AD risk by demonstrating a significantly higher burden of CHIP mutations in the blood of AD patients, particularly those with the *APOE* ε3/ε3 genotype. This association was robustly supported by two different sequencing methods and in two independent cohorts. Previous studies have reported conflicting findings, suggesting that CHIP mutations have no effect^13,14^ or even a protective effect^15^ on AD. These discrepancies may be attributed to methodological differences. Specifically, prior studies used conventional WES or WGS data with lower depth, limiting their ability to detect low VAF mutations. In contrast, our study employed deep panel sequencing, enabling the detection of low-VAF mutations. Given that the most prominent differences in mutation burden between AD cases and controls were observed in these low-VAF mutations, some prior studies may have been unable to capture these critical distinctions, though the large scale of the ADSP captures a risk effect even at lower WGS coverage. Furthermore, we observed while doing this study that age was not well matched between AD cases and controls in the ADSP WES dataset (Supplementary Table 1 and 2), which was analyzed as part of one previous study^15^. Since CHIP mutations accumulate nonlinearly with age, any age imbalance could contribute to inconsistent findings, even after trying to control for it post hoc. A more detailed discussion of these discrepancies is provided in the Supplementary Note.

Our findings, along with reports of increased somatic CHIP mutations in AD brains^10^, suggest that CHIP mutations may play an important role in AD pathogenesis. Hematopoietic cells can migrate to the brain and differentiate into macrophages or microglia-like cells, especially during aging or neurodegenerative processes when blood-brain barrier integrity is often compromised^19,20^. Previous studies have shown that blood-derived macrophages carrying the *APOE* ε4 allele exhibit phagocytic inefficiency and dysfunctional inflammation^21^, a phenotype also observed in macrophages harboring CHIP mutations^17,22^. This may explain observed differences in CHIP mutation burden across *APOE* genotypes in our study, wherein only AD patients with the *APOE* ε3/ε3 genotype, but not the ε3/ε4 genotype, exhibit increased CHIP mutations, because both the *APOE* ε4 allele and CHIP mutations could independently contribute to AD pathogenesis by impairing the clearance of amyloid-beta and promoting a neuroinflammatory environment that ultimately leads to neuronal death. Our study highlights that somatic CHIP mutation is a new genetic risk factor for AD, potentially contributing to neurodegeneration through a mechanism distinct from the established *APOE* ε4 allele.

## Methods

### Sample information

All participants subject to targeted panel sequencing were of Korean ethnicity and had participated in previous studies^1,2^. Peripheral blood samples were collected from all participants at Samsung Medical Center (Seoul, South Korea). Genomic DNA was extracted from peripheral blood cells using standard procedures. All sequencing procedures were performed at Samsung Medical Center in three sequencing batches. Samples were excluded based on the following criteria: 1) inaccurate age information, 2) low sequencing yields, and 3) evidence of oxidative damage during sample preservation^3–6^. Specifically, samples were excluded for oxidative damage if they met the following criteria: (a) more than half of their somatic mutations were C>A substitutions, or (b) the cosine similarity of their trinucleotide mutational spectrum to the spectrum of samples meeting criterion (a) was greater than 0.25. In total, 298 participants were included in our somatic mutation analysis (Extended Data Table 1), comprising 108 Alzheimer’s disease (AD) patients and 190 neurotypical controls, including 154 age-matched individuals and 36 centenarians. This study was conducted in accordance with all relevant ethical regulations and the principles of the Declaration of Helsinki. The study protocols were approved by the Institutional Review Boards (IRB) at the Samsung Medical Center (Seoul, South Korea) under approval numbers SMC IRB 2014-10-078 (for AD and control samples) and SMC IRB 2023-05-112 (for centenarian samples). A waiver of informed consent was granted by the IRB for all participants included in this study.

For AD patients, cognitive function was assessed using the Mini-Mental State Examination (MMSE). The MMSE is a widely used test for assessing cognitive impairment, with scores ranging from 0 to 30. In our AD cohort, MMSE scores ranged from 0 to 26, encompassing a spectrum of cognitive impairment from severe to mild. We found no significant association between MMSE scores and either the presence or burden of somatic mutations involved in clonal hematopoiesis of indeterminate potential (CHIP) in this cohort, even when considering *APOE* genotype (Extended Data Fig. 5).

### Panel design, sequencing, and somatic mutation calling

Targeted sequencing was performed using a panel of 149 genes previously described by Huang et al.^7^. This panel includes frequently mutated oncogenes and tumor suppressor genes associated with various cancers and clonal hematopoiesis. Library preparation was performed using the SureSelect XT HS2 DNA Reagent Kit (Agilent Technologies). Sequencing was conducted on Illumina NovaSeq 6000 platforms with 150 bp paired-end reads.

Somatic mutation calling was performed using the “sensitive” pipeline described in Huang et al.^7^. Briefly, this pipeline uses unique molecular identifiers (UMIs) to group reads originating from the same DNA molecule, generating consensus reads with increased accuracy. The UMI information of each read was first extracted from the fastq files by AGeNT’s Trimmer (v2.0.2). Reads were aligned to the GRCh37 human reference genome using BWA-MEM(v0.7.15)^8^, and consensus reads were generated using AGeNT LocatIt (v2.0.2). Somatic single nucleotide variants (sSNVs) and indels (sIndels) were called using MosaicHunter (v1.0)^9^ and Pisces (v5.3)^10^, respectively. A series of filters were applied to remove germline variants and sequencing artifacts. The “sensitive” pipeline allows for the detection of low-frequency variants that are either exclusive to or enriched in the AD or control group.

### Functional annotation of somatic mutations

We used ANNOVAR (v2015MAR22)^11^ to annotate somatic mutations based on their genomic location, classifying them into the following categories; 5’ UTR, exonic, 3’ UTR, splicing, and intronic. Exonic mutations were further classified based on their predicted impact on amino acid sequence, including nonsynonymous SNV, synonymous SNV, stopgain SNV, stoploss SNV, frameshift insertion, frameshift deletion, nonframeshift insertion, nonframeshift deletion.

We further classified a subset of somatic mutations as CHIP mutation based on the comprehensive CHIP mutation table curated by Bouzid et al.^12^. This table includes locus information for CHIP mutations across 73 genes known to be recurrently mutated in clonal hematopoiesis. Somatic variants identified in our study were compared against this table, and those matching the defined loci were annotated as CHIP mutations.

### Amplicon sequencing

Amplicon sequencing was performed to validate candidate somatic mutations and estimate mutant allele fractions. We selected 47 variants for validation which spans a range of variant allele fraction (VAF), including 22 variants annotated as CHIP. For each variant, two or three primer pairs of amplicons were designed. PCR primers were synthesized through standard oligo synthesis, and conventional PCR was conducted. After PCR, electrophoresis was used to verify the bands. The resulting PCR product DNA was adjusted to the desired concentration according to the NGS protocol to prepare the library, omitting the fragmentation process. Library preparation was performed using the TruSeq DNA PCR-Free Library Prep Kit (Illumina, Inc.). Sequencing was conducted on Illumina NovaSeq 6000 SP platforms with 150 bp paired-end reads.

Sequencing reads were aligned to the GRCh37 human reference genome using BWA-MEM (v0.7.15)^8^ and processed with GATK (v3.6) for indel realignment^13^. sSNV calls were validated based on the read fraction of the mutant allele, requiring the mutant allele read count to be greater than the read count of any error alleles at the same position. sIndel calls were manually verified using Integrative Genomics Viewer (v2.4.10)^14^. All variants analyzed by amplicon sequencing were classified into three validation categories: 1) Somatic-I: All amplicons targeting the variant were validated. 2) Somatic-II: At least one amplicon targeting the variant was validated. 3) FP (False Positive): None of the amplicons were validated.

### Mutation burden analysis

Somatic mutation burdens were calculated by dividing the total number of somatic mutations by the total size of adequately powered genomic regions with >200X coverage for the panel sequencing data. To compare mutation burdens between AD and control groups, we used the odds ratio and the two-sample Z-test of proportion.

Logistic regression analysis was performed to assess the association between somatic mutation burden and AD status while controlling for potential confounding factors. AD status was modeled as the dependent variable, and the count of somatic mutations, age, sex, and *APOE* genotype were included as independent variables.

### dN/dS analysis

To assess the selective pressures acting on somatic mutations in AD, we performed dN/dS analysis comparing the rate of protein-altering substitutions (dN) to that of synonymous substitutions (dS). A dN/dS ratio greater than 1 indicates positive selection, suggesting that the mutations are being favored by natural selection, whereas a dN/dS ratio less than 1 indicates negative selection.

We calculated dN/dS ratios for all genes in our targeted panel, as well as for the top six most frequently mutated genes. To account for the different mutation rates across genes and genomic regions, we used the dNdScv R package^15^, which employs a maximum-likelihood approach to estimate dN/dS while correcting for these biases.

### ADSP analysis

To determine the CHIP status of ADSP participants, we restricted our analysis to participants whose DNA was extracted from blood and annotated all variants present on genes listed as CHIP-related by Bouzid et al (2023)^12^. All variants and consequences listed in their Supplementary Table 1 were considered, and we computed the VAF for each individual at these variants’ positions. While Bouzid et al.^12^ list is comprehensive, it is approximative for some genes listing any variant with a given consequence as CHIP-related, our data suggest that some of these are germline, to this aim, we selected variants with median VAF across carriers (individuals with at least one alternate count) below 40%. In ADSP WGS data (30x coverage), to remove artifacts, we excluded variants with filter != PASS; variants with median_read_depth×median_VAF < 3 (ensuring that for 50% carriers at least 2 reads support the CHIP status); variants present in more than 36 participants as a given CHIP variant should not be seen in more than 0.5% of the population^16^. CHIP participants in WGS were thus defined as individuals with VAF > 2% and read depth > 15 at any of the variants from Bouzid et al. (2023)^12^ that passed the filtering step listed above. In ADSP WES, due to the higher coverage (median ∼50-70x, depending on exons), we solely required variants to be present in less than 0.1% of the population^16^ (accounting for the higher age than the ADSP WGS), in addition to the VAF < 40%. The odds ratio in ADSP WES ε3/ε3 European participants (Supplementary Table 2B) was similar to the one obtained by Bouzid et al. in the same cohort, making us confident in implementing an equivalent CHIP status identification pipeline.

In ADSP WGS, we considered unique samples of 11,565 Alzheimer’s cases and 16,873 healthy controls and run a generalized linear model (Binomial family) with Python 3.9, statsmodels package testing the association of CHIP status with disease status. Genetic sex, age-at-visit, *APOE* ε2 and *APOE* ε4 dosages, and 4 principal components accounting for genetic ancestry, were included as covariates in the model. The association of CHIP with AD was also tested by *APOE* genotype, and in these stratified analyses, *APOE* ε2 and *APOE* ε4 dosages were not adjusted.

## Data availability

The panel sequencing data generated in this study will be deposited to NIAGADS (https://www.niagads.org/), with controlled use conditions set by human privacy regulations.

## Code availability

Custom bash and R scripts used in this study will be publicly available at Github upon publication.

## Acknowledgements

The authors gratefully acknowledge Boxun Zhao for helpful discussions and advice regarding library-related aspects of this project. This work was supported by R01 AG088082 (A.Y.H.); R56 AG079857 (A.Y.H., C.A.W., E.A.L.); Alzheimer’s Association Research Fellowship (A.Y.H.); Suh Kyungbae Foundation (E.A.L.); DP2 AG072437 (E.A.L.); R01 AG070921 and AG078929 (C.A.W., E.A.L.); Allen Discovery Center program, a Paul G. Allen Frontiers Group advised program of the Paul G. Allen Family Foundation (C.A.W., E.A.L.); NIH funding source of Stanford’s Center for Clinical and Translational Education and Research award, under the Biostatistics, Epidemiology and Research Design (BERD) Program: 1UM1TR004921-01 (Y.L.G.); Korea Dementia Research Project RS-2024-00339665 through the Korea Dementia Research Center (KDRC), funded by the Ministry of Health & Welfare and Ministry of Science and ICT, Republic of Korea (J.K.); PRMRP Discovery Award W81XWH2010028 (Z.Z.); the Edward R. and Anne G. Lefler Center postdoctoral fellowship (Z.Z.); the American Heart Association Career Development Award 23CDA1046074 (Z.Z.).

## Author Contributions

E.A.L. conceived the study. J.C., K.S.P., Y.L.G., J.K., A.Y.H., and E.A.L. designed the experiments and analyses. K.S.P., J.P., Y.C., and J.K. performed sample collection, processing (including storage and preservation), and sequencing. Z.Z., L.E., and I.R. designed and validated sequencing probes for panel sequencing and amplicon sequencing. J.C. and A.Y.H. performed variant calling and conducted statistical analyses. Y.L.G. performed statistical analyses on ADSP data. C.A.W. helped interpret the results. J.C. drafted the initial manuscript, with significant contributions from A.Y.H., K.S.P., and Y.L.G. A.Y.H. and E.A.L. supervised the study. All the authors read, revised and approved the manuscript.

---

**Extended Data Figure 1.**
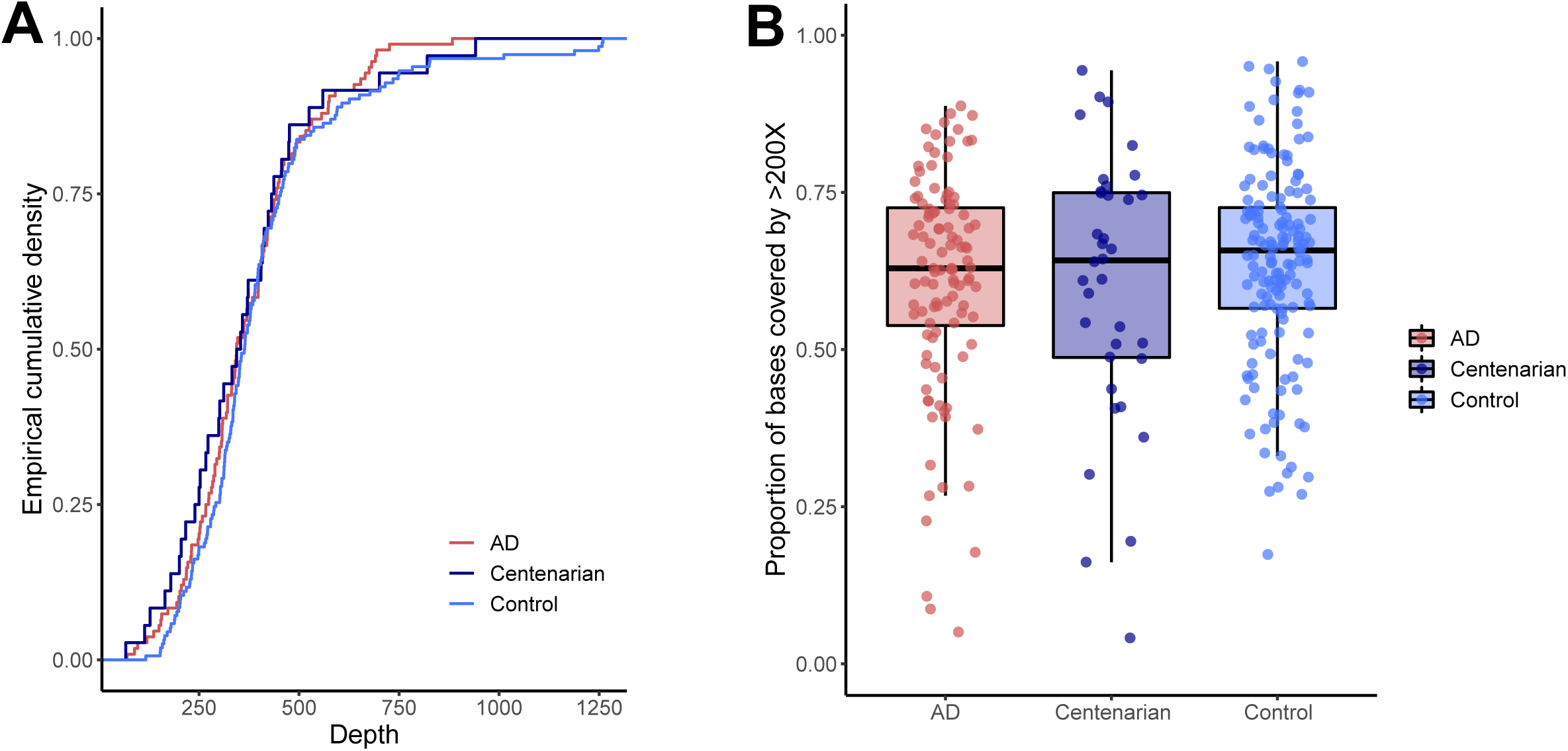
Comparison of sequencing depth and coverage across different sample groups of targeted panel sequencing data. **A.** Cumulative density plots illustrating the distribution of sequencing depths across samples. The x-axis represents sequencing depth, and the y-axis represents the empirical cumulative density reflecting the cumulative proportion of samples sequenced at or below a given depth in the AD, centenarian, and control groups. **B.** Boxplots showing the proportion of genomic bases covered by more than 200X depth for each sample group. In the boxplots, the center line indicates the median value. The upper and lower limits of the box represent the 75th and 25th percentiles (the interquartile range, IQR), respectively. The whiskers extend to 1.5 times the IQR from the box limits, or to the most extreme data point within that range if no points are further out. Individual data points are overlaid, jittered for visibility. No significant differences were observed in sequencing depth or coverage among the three groups.

**Extended Data Figure 2.**
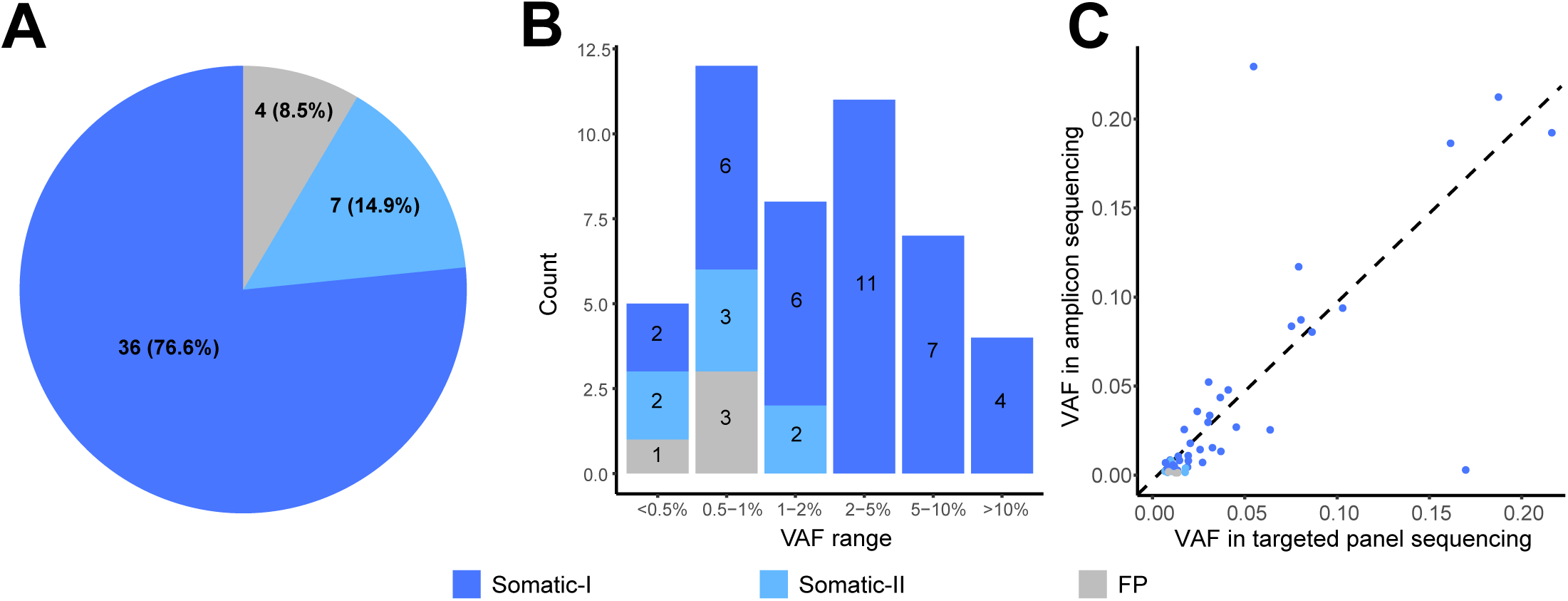
Validation of somatic mutations using amplicon sequencing. **A.** Pie chart summarizing the validation results of 47 selected somatic mutations. Variants are categorized as “Somatic-I” (validated in all designed amplicons, 36 variants, 76.6%), “Somatic-II” (validated in at least one amplicon, 7 variants, 14.9%), or “False Positive” (FP, 4 variants, 8.5%). **B.** Bar plot showing the distribution of variant allele fractions (VAFs) for all variants analyzed by amplicon sequencing, grouped by validation category. The numbers on each bar represent the count of each validation group in each VAF group. **C.** Scatter plot demonstrating the strong concordance of VAF between targeted panel sequencing and amplicon sequencing. The dashed line denotes a theoretical 1:1 correlation.

**Extended Data Figure 3.**
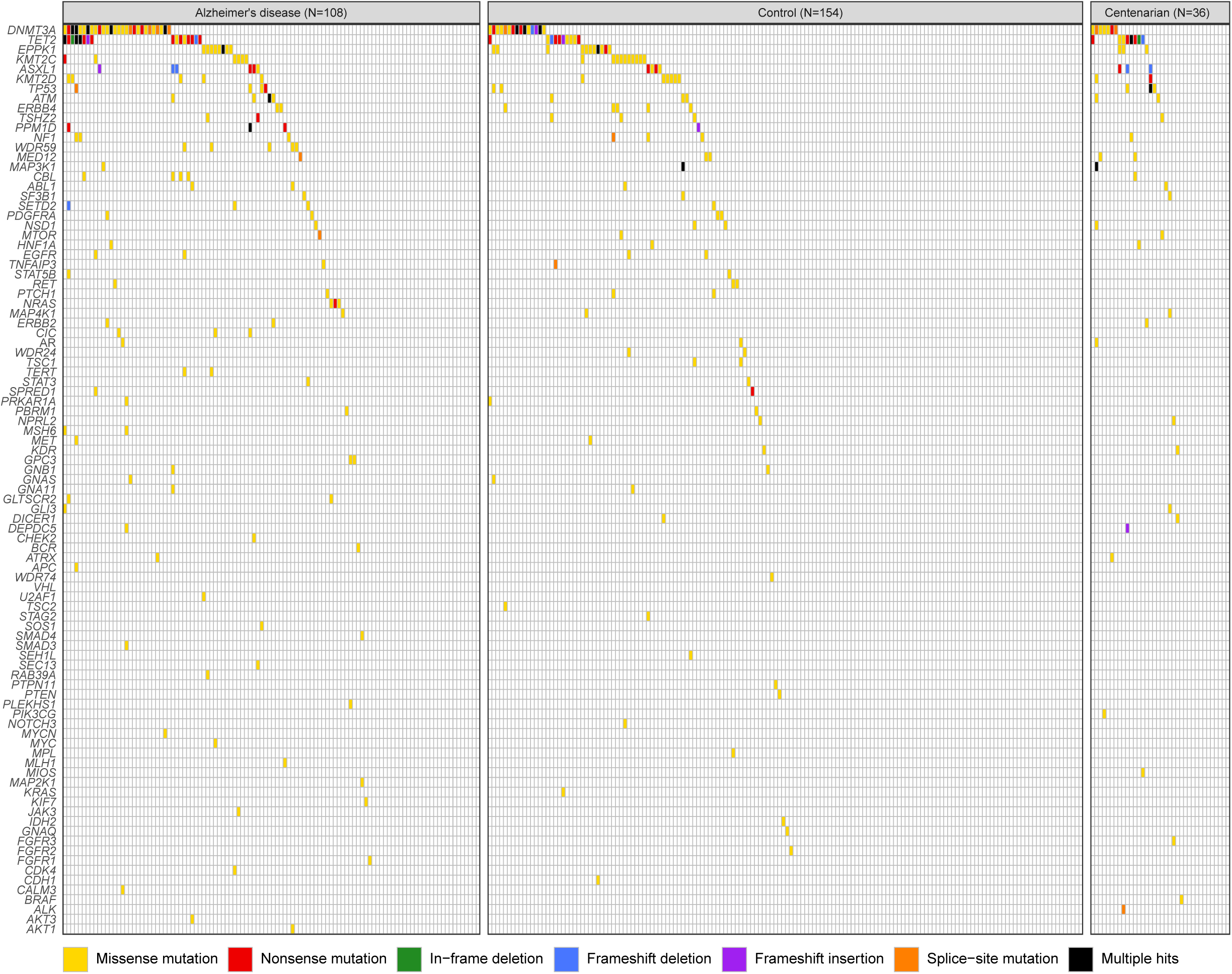
Gene-wide landscape of all non-silent exonic mutations across AD, age-matched control, and centenarian groups. The colors represent different types of mutation: missense SNV, nonsense SNV, in-frame deletion, frameshift deletion, frameshift insertion, splice-site mutation, and multiple hits. Overall, a higher proportion of AD patients carried somatic mutations among these cancer driver genes compared to age-matched controls.

**Extended Data Figure 4.**
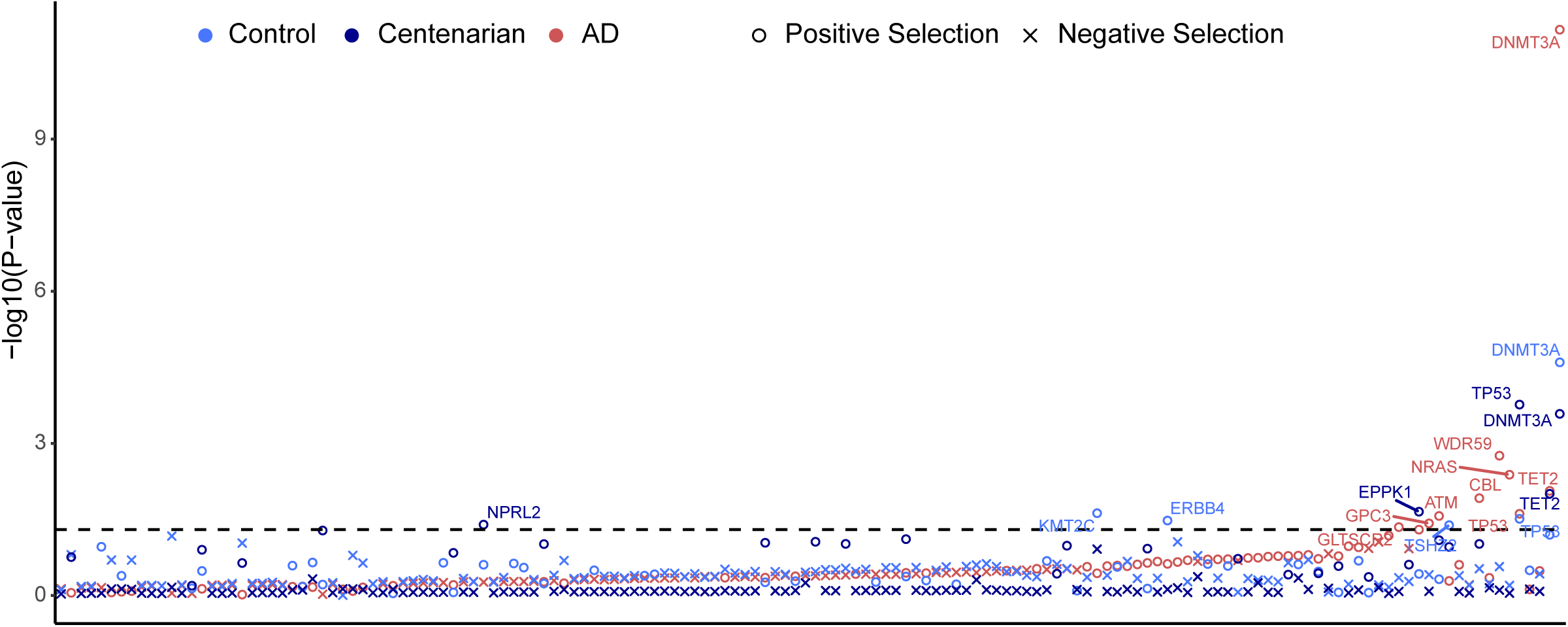
dN/dS analysis indicating positive selection in certain genes across AD, control, and centenarian groups. The x-axis denotes each of the 149 cancer driver mutations in our panel sequencing, and the y-axis denotes the log10-transformed p-value, indicating the statistical significance of dN/dS ratio deviations from the neutral expectation for each gene. Genes with signal of positive selection are indicated with an open circle, while those with signal of negative selection are marked with an “x”. The horizontal dashed line represents the significance threshold (p = 0.05). Gene names are labeled for genes showing significant (p < 0.05) dN/dS ratios.

**Extended Data Figure 5.**
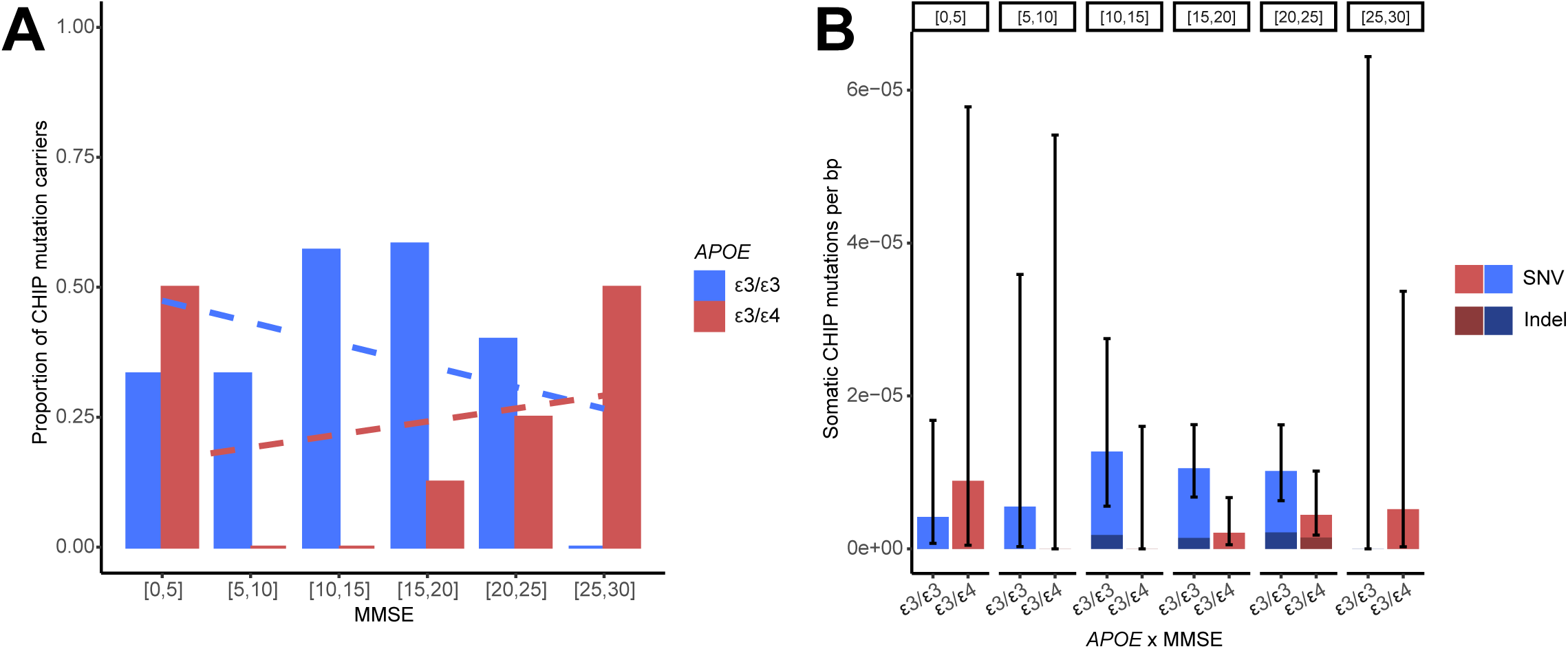
Association between CHIP mutations and Mini-Mental State Examination (MMSE) scores stratified by *APOE* genotype in AD samples. **A.** Proportion of CHIP mutation carriers across different MMSE score ranges, stratified by *APOE* ε3/ε3 and ε3/ε4 genotypes. Dashed lines represent the trend lines for each genotype, illustrating the overall trend in the proportion of CHIP carriers across MMSE ranges. **B.** Number of somatic CHIP mutations per bp for SNVs and indels across different *APOE* genotype and MMSE score combinations. Error bars represent 95% CIs. No significant correlation was observed between MMSE scores and the presence or burden of CHIP mutations within either *APOE* genotype group.

