## Supplementary Note for "Clonal Hematopoiesis Mutations Increase Risk of Alzheimer’s Disease with *APOE* ε3/ε3 Genotype"

**Sequencing depth, quality control, and comparison to previous studies**

This study utilized targeted sequencing of a 149-gene panel to investigate somatic mutations, including those associated with clonal hematopoiesis of indeterminate potential (CHIP), in peripheral blood samples from Alzheimer's disease (AD) patients, age-matched controls, and centenarians. To ensure high-quality data and reliable variant calling, we implemented multiple quality control measures and achieved high average sequencing depth (~400X) after generating consensus sequences using molecular barcoding. Sequencing depth and coverage were comparable between AD and control groups (Extended Data Fig. 1), minimizing potential bias from technical variation.

Our study achieved significantly higher sequencing depth than previous studies that investigated CHIP mutations in AD, which relied on whole-exome sequencing (WES; ~50-70X in ADSP WES samples analyzed by Bouzid et al.[1]) or whole-genome sequencing (WGS; ~30X in TOPMed samples analyzed by Bouzid et al.[1]). The use of targeted sequencing in our study allowed us to achieve much higher depth concentrated in regions of interest, enhancing sensitivity for detecting low-fraction somatic mutations (particularly those with VAF < 5%), which show the strongest signal of elevated mutation burden in AD (Fig. 1F and G); such mutations would be largely undetectable in WES or WGS at conventional depths. Additionally, The incorporation of molecular barcoding mitigated sequencing artifacts from PCR amplification and base-calling, improving the accuracy of somatic mutation detection—particularly for low-fraction ones—and achieving an 91.5% validation rate (Extended Data Fig. 2), as previously demonstrated by Huang et al.[2].

**Addressing potential confounding effects of age in previous studies and data selection for the current study**

It has long been known that CHIP prevalence increases with age, and this increase becomes quadratic after approximately 70 years of age[3-5]; therefore, studies examining the relationship between CHIP and aging-related clinical outcomes, such as AD, must carefully ensure age-matching between cases and controls to enable unbiased comparisons.

A previous study by Bouzid et al.[1] investigated the association between CHIP and AD and analyzed the first release of ADSP WES dataset[6], comprising approximately 10,000 samples. However, the study design of the ADSP WES cohort resulted in an enrichment of younger-onset cases and older controls, with controls being, on average, 10 years older than cases[7] (Supplementary Table 1). Therefore, their findings regarding the protective effect of CHIP against AD could be confounded by this age difference, as the higher burden of CHIP in their control groups could be driven by age rather than a true protective effect against AD.

To mitigate this potential confounding effect, we analyzed the ADSP WGS dataset (approximately 36,000 samples, ng00067.v12 release), which includes a larger number of samples and, importantly, exhibits improved, more balanced age-matching between cases and controls (Supplementary Table 1). While the precise age at blood draw for genomic sequencing is not available in the ADSP and NIAGADS databases, the age at study enrollment provides a reasonable proxy. By leveraging this dataset, we identified a significantly elevated burden of CHIP mutations in AD compared to age-matched controls (Fig. 1I), consistent with our findings from targeted panel sequencing. This rigorous consideration of age-matching in both panel sequencing and ADSP WGS strengthens the robustness of our findings and enables a more precise assessment of the true relationship between CHIP and AD risk.
